# *ATP1A1*-linked diseases require a malfunctioning protein product from one allele

**DOI:** 10.1101/2023.03.05.531165

**Authors:** Kerri Spontarelli, Victoria C. Young, Ryan Sweazey, Alexandria Padro, Jeannie Lee, Tulio Bueso, Roberto M. Hernandez, Jongyeol Kim, Alexander Katz, Francis Rossignol, Clesson Turner, Caralynn M. Wilczewski, George L. Maxwell, Miguel Holmgren, Jeremy D. Bailoo, Sho T. Yano, Pablo Artigas

## Abstract

Heterozygous germline variants in *ATP1A1*, the gene encoding the α1 subunit of the Na^+^/K^+^-ATPase (NKA), have been linked to diseases including primary hyperaldosteronism and the peripheral neuropathy Charcot-Marie-Tooth disease (CMT). *ATP1A1* variants that cause CMT induce loss-of-function of NKA. This heterodimeric (αβ) enzyme hydrolyzes ATP to establish transmembrane electrochemical gradients of Na^+^ and K^+^ that are essential for electrical signaling and cell survival. Of the 4 catalytic subunit isoforms, α1 is ubiquitously expressed and is the predominant paralog in peripheral axons. Human population sequencing datasets indicate strong negative selection against both missense and protein-null *ATP1A1* variants. To test whether haploinsufficiency generated by heterozygous protein-null alleles are sufficient to cause disease, we tested the neuromuscular characteristics of heterozygous *Atp1a1*^+/-^ knockout mice and their wildtype littermates, while also evaluating if exercise increased CMT penetrance. We found that *Atp1a1*^+/-^ mice were phenotypically normal up to 18 months of age. Consistent with the observations in mice, we report clinical phenotyping of a healthy adult human who lacks any clinical features of known *ATP1A1*-related diseases despite carrying a protein-null early truncation variant, p.Y148*. Taken together, these results suggest that a malfunctioning gene product is required for disease induction by *ATP1A1* variants and that if any pathology is associated with protein-null variants, they may display low penetrance or high age of onset.

## Introduction

Mutations of the Na^+^/K^+^ pump α1 subunit gene (the human gene will be referred to as *ATP1A1* and the mouse gene as *Atp1a1*) cause various disorders, including hyperaldosteronism with secondary hypertension affecting the endocrine system (*1-3*), Charcot-Marie-Tooth disease (CMT, *4, 5*) and hereditary spastic paraplegia (*6*), affecting the neuromuscular system, and hypomagnesemia with seizures and intellectual disability (*7*), disturbing the renal and central nervous systems (reviewed by *8*).

The Na^+^/K^+^ pump or Na^+^,K^+^-ATPase (NKA) hydrolyzes ATP to establish electrochemical gradients for Na^+^ and K^+^ across the plasma membrane of all human cells. These gradients are essential for cellular survival, as they drive electrical signaling and secondary-active transport. Different tissues express distinct NKA isozymes formed by association of one α-subunit isoform (α1-α4) with one β-subunit isoform (β1-β3). NKAs formed by the α1 subunit are ubiquitous, while α2 pumps predominate in skeletal muscle and glia, and α3 pumps are expressed in neurons. Notably, sensory and motor neurons express both α1 and α3 isoforms, with α1 predominating in axons (*4*), which may explain why α1 disease-associated variants predominantly cause CMT type 2 in which axonal degeneration leads to progressive strength and sensory loss starting from the distal limbs, although an *ATP1A1* variant causing demyelinating CMT was recently described (*9*).

Heterozygous loss-of-function, with a missense variant that partially reduces Na^+^/K^+^-transport by the gene product of one allele, has been proposed to cause some *ATP1A1* disease phenotypes. Electrophysiological studies in *Xenopus* oocytes and adrenal NCI-H295R cells expressing disease-related NKA α1 variants reported depolarizing inward currents through most (but not all) variants linked to hyperaldosteronism (*2, 8, 10, 11*) or hypomagnesemia with refractory seizures (*7, 12*), providing a plausible gain-of-function mechanism for these disease-associated variants. However, variants causing neuropathies seem to lack such inward “leak” currents (*4-7, 9-11*). Moreover, functionally characterized CMT variants impair NKA function to distinct degrees. Families carrying mutations found to be the most deleterious in functional studies have higher penetrance and disease severity, while the family carrying L48R, the variant with the mildest functional effect, has incomplete CMT penetrance with variable age of disease onset (8-50 years old, *4*).

Therefore, we asked whether heterozygous protein-null alleles are also sufficient to cause disease. Sequencing datasets of the human population indicate strong negative selection against both missense and protein-null variants, with low ratios of observed/expected variant frequency in gnomAD v.2.1.1: 0.29 for missense (*Z* = −6.22) and 0.04 for predicted loss-of-function (pLI=1) variants (*13*). Despite the nonzero allele frequency, the human phenotype of protein-null variants is unknown, as such individuals have yet to be reported. On the other hand, to date, the reported characteristics of heterozygous *Atp1a1*^*+/-*^ knockout mice up to 6 months of age show minimal, if any, phenotypic differences with wildtype mice (*14-16*). As CMT symptoms progress through the patient’s lifespan, and because previous studies of *Atp1a1*^+/-^ mice did <u>not</u> exhaustively tested neuromuscular characteristics, we compared the neuromotor characteristics of heterozygous *Atp1a1*^*+/-*^ knockout mice with wildtype littermates, up to 18 months old. Moreover, we evaluated whether a long-term exercise regime increases CMT penetrance by increasing the Na^+^ load in nerves with half the gene product.

Somewhat surprisingly, given the low ratio of human protein-null variants reported, we found that *Atp1a1*^*+/-*^ mice were phenotypically normal throughout their lifetime, and that exercise did not alter CMT penetrance. Consistent with these results we present the clinical evaluation of a healthy adult female volunteer carrying the early truncation variant Y148*, which was identified in a next-generation sequencing screen and had a protein-null effect *in vitro*. Taken together, these results suggest that a malfunctioning gene product is required for disease induction by *ATP1A1* variants and that protein-null variants do not cause highly penetrant *ATP1A1*-related diseases.

## RESULTS

We confirmed the reduction in α1 protein at the axolemma from sciatic nerves of *Atp1a1*^*+/-*^ mice by Western blotting (Fig. 1). Sciatic nerves from five-month-old mice from each genotype were used for the membrane preparation. Each lane in Fig. 1A was loaded with the indicated protein concentration and probed with rabbit anti α1 (*top*) and mouse anti α3 antibodies (*bottom*). The plot of the band fluorescence intensity vs. loaded protein (Figure 1B) follows a linear dependence (below 7 μg of protein for the α1 antibody). The ∼50% reduction in α1 observed in *Atp1a1*^*+/-*^ heterozygous mice compared to wildtype littermates is <u>not</u> accompanied by a detectable increase in α3 or α2, with the latter being undetectable in axolemma membrane preparation (Figure 1C).

**Figure 1.**
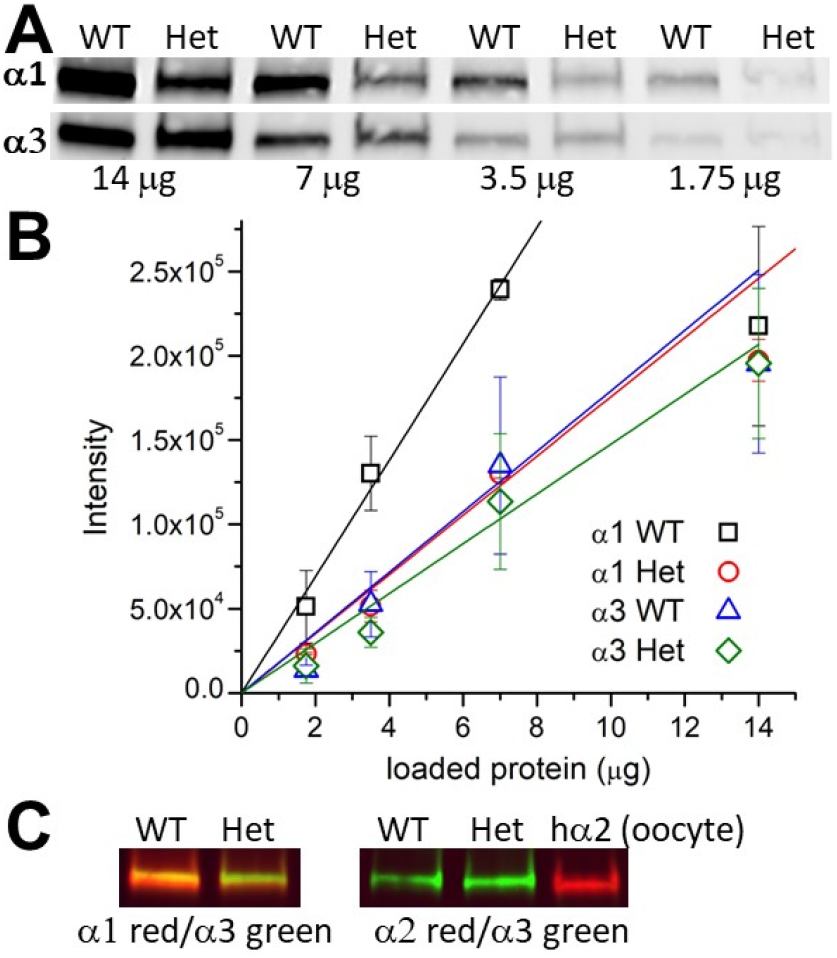
Nerves from *Atp1a1*^*+/-*^ mice have 50% lower α1 protein levels. **A)** Western blot (WB) of sciatic nerve axolemma protein preparation of WT and *Atp1a1*^*+/-*^ (Het) sciatic nerves (five WT and six *Atp1a1*^*+/-*^ 5-month-old littermates were pooled in each). Different amounts of protein were loaded, as indicated. The WB was probed with the anti-α1 NASE antibody (rabbit) and the anti-α3 XVIF9-G10 (mouse) antibody. Goat anti-rabbit and goat anti-mouse secondaries (Licor) emitted, respectively at 680 and 800 nm. **B)** Mean fluorescence intensity from two WB (in arbitrary units, AU) as a function of the protein loaded in the well. Line plots are linear regressions fitted to the data up to 7 μg (after which the α1 signal saturates), with slopes 17,916 AU/μg protein (α3 WT blue triangles), 36,991 AU/μg (α1 WT, black squares), 14,756 AU/μg (α3 Het green diamonds) and 18,288 (α1 Het red circles). **C)** WB where the membrane was cut in two to probe with different rabbit-raised isoform-specific antibodies; NASE against α1 (*left*) and HERED against α2 (*right*). As indicated, the lanes were loaded with WT or *Atp1a1*^+/-^ axolemma protein (8 μg) and with a membrane prep from oocytes expressing human α2, as a control for HERED. The composite orange and yellow color indicate the presence of both red and green signals in the membrane prep from mice stained with α1 and α3 antibodies. The green band when staining the mouse membranes for α2 and α3 indicates that the nerve membranes lack α2.

### Cross-sectional neuromotor evaluation of Atp1a1^+/-^ mice

We performed a battery of tests for the neuromotor evaluation of *Atp1a1*^*+/-*^ and their wildtype littermates from 1 to 18 months of age (Figures 2-3, Table 1). Balance, strength, and motor coordination was measured using the balance beam, vertical pole, and rotarod tests, respectively. The primary outcome variables measured in each test are those affected in neuropathic mice: the latency to cross in the balance beam (*17, 18*) (Figure 2A), the latency to descend in the pole test (*19, 20*) (Figure 2B), and the latency to fall off the accelerating rotarod (*21, 22*) (Figure 2C). In general, the data show that *Atp1a1*^*+/-*^ mice lack neuropathic characteristics (Table 1, Figure 2 A-C). Notably, however, two *Atp1a1*^*+/-*^ mice took much longer to cross the balance beam at 12 and 18 months (Figure 2A). These mice slid across the bar, using the sides of the bar to push themselves forward, instead of walking on top of the bar. Such sliding motor behavior was infrequent and inconsistent within and across trials in other mice at all ages. Specifically, two *Atp1a1*^*+/-*^ mice switched between sliding and walking at 12 months (of 39 mice total), which progressed to sliding only at 18 months (of 33 mice total); four additional *Atp1a1*^*+/-*^ mice slid/walked, one wildtype mouse slid/walked, and two wildtype mice slid at 18 months.

**Figure 2.**
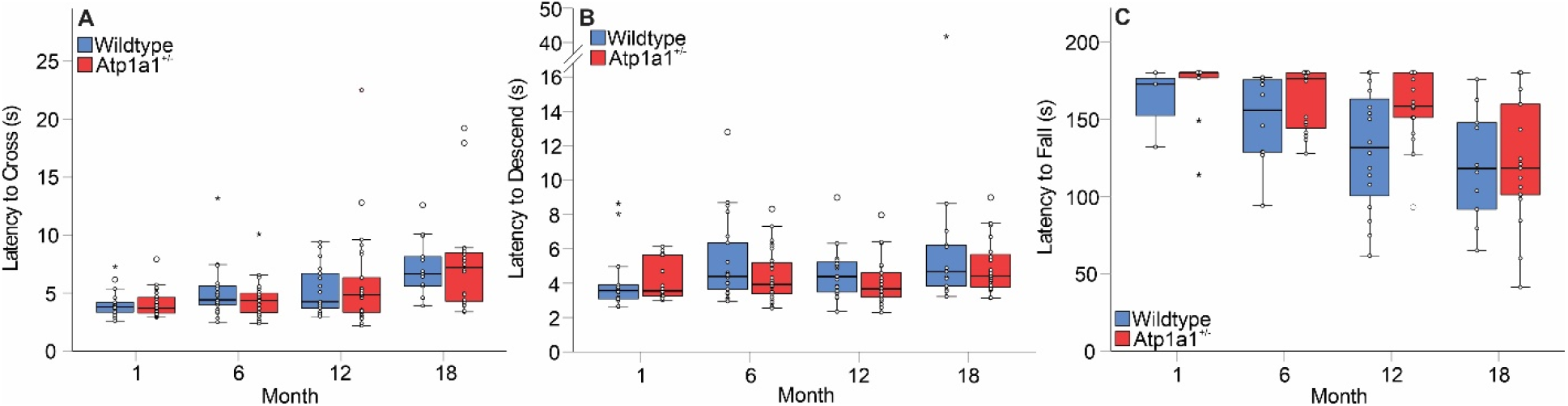
Distribution of outcomes by genotype across month in the balance beam (**A**), pole test (**B**), and rotarod (**C**). Extreme values (outside the first quartile) are indicated by wide circles while highly extreme values (outside two quartiles) are designated by stars. The time to fall in the rotarod at 12 months of age was the only statistically significant difference observed between the two genotypes.

**Figure 3.**
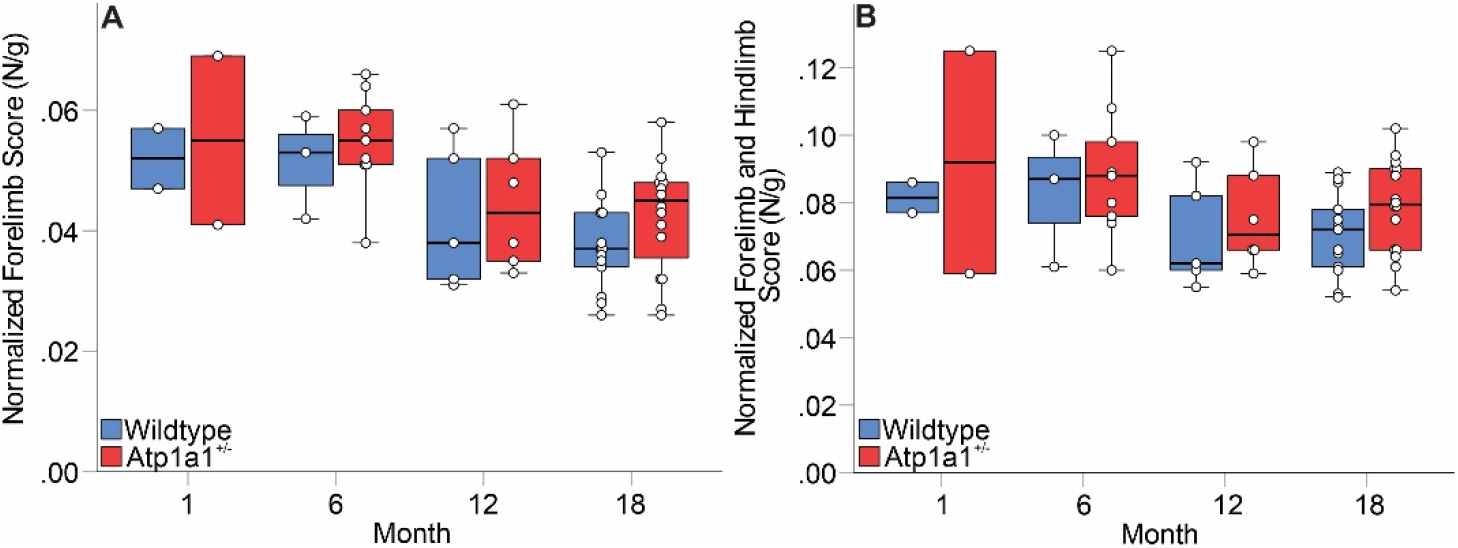
Grip strength is unaltered in *Atp1a1*^*+/-*^ mice. **A)** Forelimb strength (only front legs allowed to grip), **B**) forelimb and hindlimb strength (with all 4 legs grasping the grid simultaneously). Force was normalized to body weight. The lower number of data points in younger animals reflects addition of this test later in the study.

**Table 1.** Summary of Mann-Whitney U statistics comparing neuromotor performance of the two genotypes. All but one value (at 12 months, highlighted in red) were <u>not</u> statistically different (p < 0.05).

| Test | Outcome Measure | Age |  |  |  |
| --- | --- | --- | --- | --- | --- |
|  |  | 1 | 6 | 12 | 18 |
| Balance Beam | *Time to Cross* | -0.47 | 0.15 | 0.07 | 0.83 |
| Pole Test | *Time to Descend* | -0.59 | 1.37 | 1.45 | 0.44 |
| Rotarod | *Time to Fall* | -0.79 | -1.92 | -2.21 | -0.20 |
| Grip Strength | *Forelimb* | 0.00 | -0.46 | -0.55 | -1.71 |
|  | *Forelimb & Hindlimb* | 0.00 | -0.28 | -0.73 | -1.84 |

When hung from the tail, wildtype mice generally splay their legs away from the center while mice with peripheral neuropathy clasp their legs towards the center of the body line. Quantification for this so-called hindlimb clasping test varies (*18, 21-23*). We also found that previously unreported methodological variables can significantly alter the results from individual mice (described in Methods). Once these were identified, mice were hung by the tip of the tail, suspended over an empty cage with a grid lid, with the forelimbs approximately 3 cm from the lid. We did <u>not</u> observe differences in the hindlimb splaying/clasping behavior of both genotypes.

Patients with CMT first lose strength in their distal muscles (hands and feet), thus impacting grip strength early in the disease (*24*). Later in our study, we measured grip strength (*21, 22, 25*) due to a lack of significant differences among genotypes in the other variables. Again, grip strength in *Atp1a1*^*+/-*^ and WT mice was statistically indistinguishable (Figure 3).

### Compound muscular action potential (CMAP)

All humans carrying the L48R disease-associated variant, including those with strength above the CMT diagnostic criteria, were reported to have reduced CMAP amplitude, indicating that asymptomatic carriers have a subclinical phenotype due to loss of nerve fibers. We tested whether aged heterozygous mice had a similar reduction of CMAP amplitude that could indicate an impairment level undetectable in the neuromotor tests. No significant differences were seen in the CMAP amplitudes of *Atp1a1*^*+/-*^ and WT mice measured at 18-months-old (Fig. 4, Table 1).

**Figure 4.**
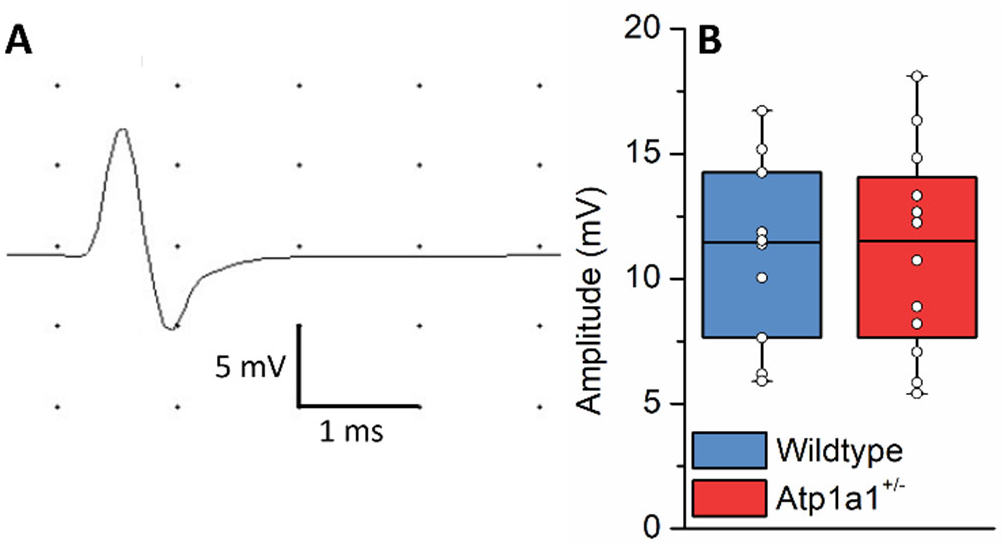
CMAP is unaltered in *Atp1a1*^+/-^ mice. **A)** Representative time-voltage trace produced by maximal stimulation of the sciatic nerve and recorded at the intrinsic muscles of the footpad of a wildtype mouse. **B)** CMAP amplitudes (average of both footpads) in wildtype (n=10) and *Atp1a1*^*+/-*^ (n=12) littermates at 18-month-old.

### Exercise does <u>not</u> increase CMT penetrance

Mice in animal facilities are sedentary (*26-29*). We reasoned that an increase in action potential propagation frequency due to regular exercise could increase the load on the reduced number of pumps in the nerves of *Atp1a1*^*+/-*^ mice, thereby increasing the likelihood of observing a CMT phenotype. A subset of 21 animals (two litters, 11 *Atp1a1*^*+/-*^ and 10 wild type) were exercised for 30 minutes at 13 m/min, 5 days a week, after the first month and until the sixth month of age (one *Atp1a1*^*+/-*^ died at 5 months old). Exercised animals were compared to non-exercised animals (28 *Atp1a1*^*+/-*^ and 20 wild type) on the same behavioral tests described above at 6 months of age. No significant differences were observed between the genotypes (Figure 5). However, exercise increased the frequency of wildtype mice reaching the 180 s ceiling, an effect that appears absent from heterozygous mice.

**Figure 5.**
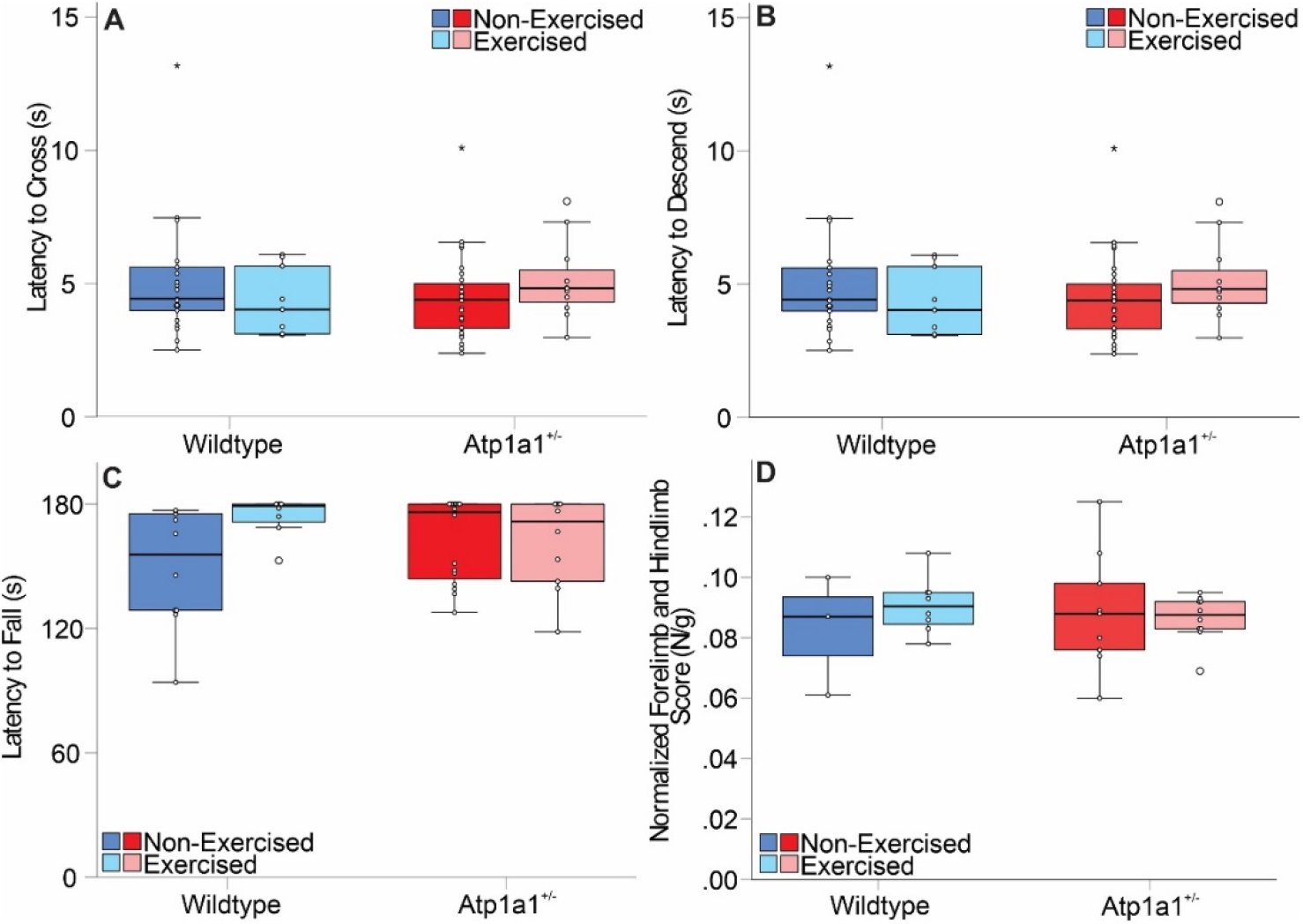
Evaluation of the effects of exercise on genotype in the distribution of outcomes at 6 months of age in the balance beam **(A)**, pole test **(B)**, rotarod **(C)** and grip strength **(D)**. Extreme values (outside the first quartile) are indicated by wide circles while highly extreme values (outside two quartiles) are designated by stars.

### Aged Atp1a1^+/-^ mice have normal serum aldosterone

Somatic mutations of *ATP1A1* in aldosterone producing adenomas cause primary aldosteronism. It was previously reported that *Atp1a1*^*+/-*^ mice had 2-fold higher aldosterone levels than wildtype mice (*16*). However, our aldosterone serum levels measured in 18 months old littermates were not significantly different, with values 232.4 pg/ml ± 75.8 (n = 8, SD) in *Atp1a1*^*+/-*^ and 191.3 pg/ml ± 56.3 (n = 9, SD) for wildtype mice, respectively.

### A healthy human volunteer lacking one ATP1A1 allele product is neurologically typical

We phenotyped a healthy 41 year old adult volunteer with a heterozygous *ATP1A1* nonsense variant predicted to lead to nonsense-mediated decay, NM_000701.7:c.444C>G; p.(Y148*). She was part of a larger genotype-driven study cohort recruited solely on the basis of *ATP1A1* variants computationally predicted to be deleterious, regardless of phenotype (manuscript in preparation). She had no history of seizures or developmental delay. There were no specific medical concerns. Features elicited on history were several lifetime migraines with visual aura; subjectively worse hearing recently; Raynaud syndrome involving painful purple blistering of all toes alleviated by warmth; one episode of pruritic hives; and bicornuate uterus. Evaluation showed normal nerve conduction and electromyography (NCS/EMG, including normal distal CMAPs), blood pressure, physical, dysmorphology and neurological exams, complete blood count, liver function testing, upright serum aldosterone (17.6 ng/dL, laboratory reference 6.5-86), plasma renin activity (1.3 ng/mL/hr, reference 0.6-3), thyroid stimulating hormone and free T4, serum and random urine electrolytes including magnesium (2.1 mg/dL serum, 1.9 mg/dL urine), and fractional excretion of magnesium (3.7%). Therefore, this individual was confirmed negative in adulthood for all known germline and somatic *ATP1A1*-related diseases (*8*), i.e. primary aldosteronism, epilepsy-renal hypomagnesemia, axonal Charcot-Marie-Tooth neuropathy, spastic paraparesis, or developmental delay.

The catalytic α subunits require β subunits to reach the plasma membrane (*30, 31*). To confirm the expectation that the Y148* protein product cannot reach the plasma membrane due to lack of β subunit interaction, we coexpressed the human β1 subunit with either human wildtype α1 or Y140*-α1 to which a YFP tag had been added in the N-terminus (Figure 6). Confocal imaging shows intense YFP fluorescence at the plasma membrane for wildtype, as expected (Figure 6A, *left*). Y148* showed variable degrees of diffuse fluorescence intensity in the cytosol (Figure 6A, *right*), but not at the plasma membrane. Western blot preparations of plasma membrane enriched preparations from transfected cells show a clear signal due to the N-terminally tagged WT pumps (∼120 kD), and recognized both with a specific α1 antibody (*32*) (Figure 6B) and a YFP antibody (Figure 6C), while Y148* was not detected with either antibody. Thus, the Y148* variant is equivalent to the lack of one allele product in the plasma membrane.

**Figure 6.**
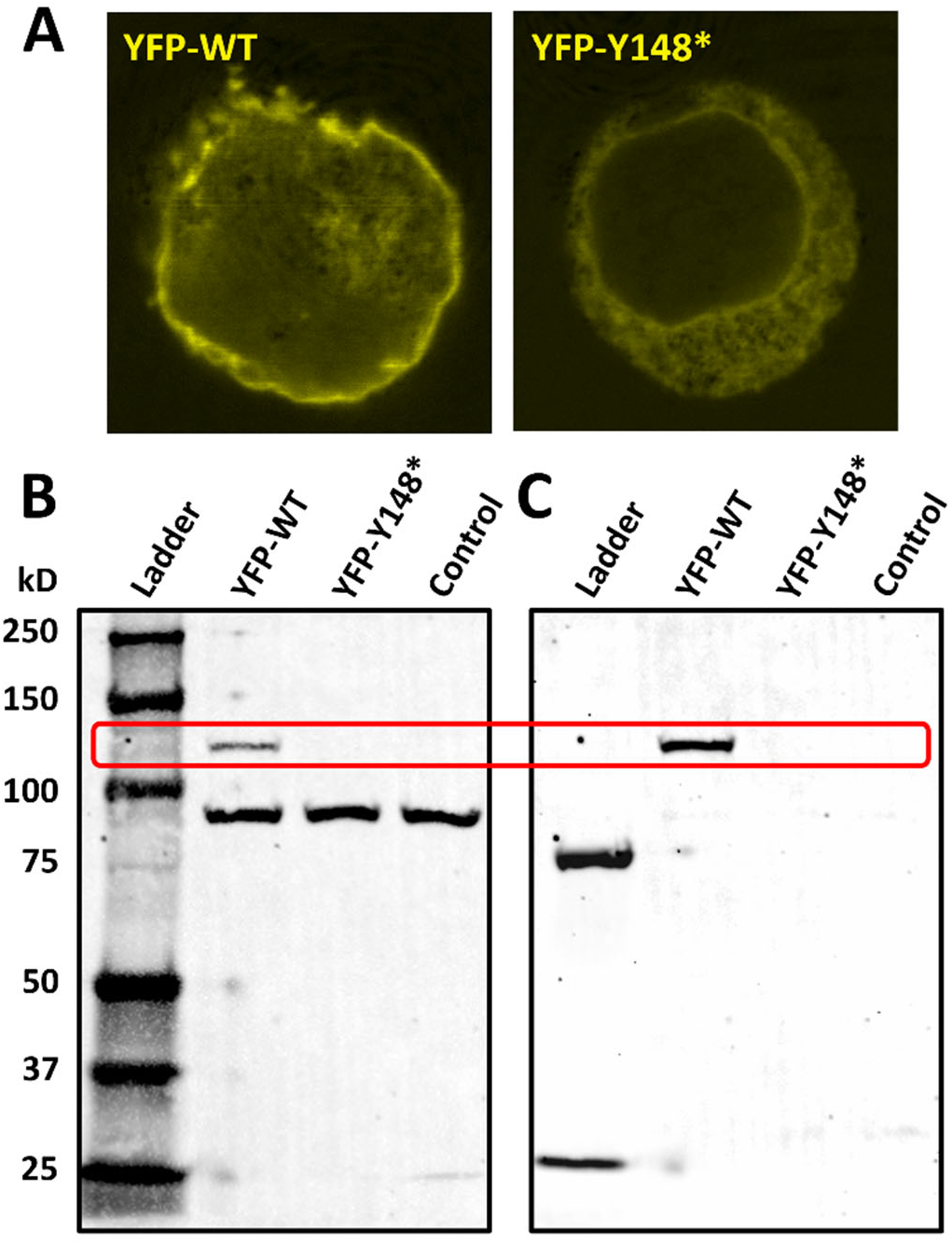
<u>Y148* is absent from plasma membrane</u>. A) Confocal image of HEK293 cells transfected with the β1 subunit and either the YFP-WT α1 subunit (*left*) or the YFP-Y148* α1 (*right*). Note high intensity of YFP fluorescence at the plasma membrane in YFP WT transfected cells, and the somewhat weaker diffuse fluorescence in the cytosol of YFP-Y148*. Images were taken on the same day with identical acquisition parameters for both transfections. **B, C)** Western blot from enriched plasma membrane preparations from HEK cells simultaneously probed with the rabbit raised anti-α1 antibody NASE (**B**) and a mouse raised anti-YFP antibody (**C**). The first lane was loaded with ladder (at the two acquisition colors the ladder indicates different weights). The other lanes were loaded with membrane preparations from HEK293 cells transfected with YFP-WT (*lane 2*), YFP-Y148* (*lane 3*) and non-transfected cells (*lane 4*). Note that while the anti-α1 NASE antibody recognizes a band near 100 kD in all three membrane loaded lanes (due to the endogenous α1 pumps) only the YFP-WT lane has an additional band at ∼ 120 kD, due to the higher molecular weight of the exogenously expressed YFP-WT. As the epitope for the anti-α1 NASE antibody (*33*) is not expected to recognize the Y148* protein, we also probed with the anti-YFP antibody, which stained a band at 120 kD corresponding to the YFP-WT construct, but the YFP-Y148* construct (expected at ∼40 kD) was not detected.

## Discussion

One of the *ATP1A1* missense variants currently known to be associated with CMT2, L48R, showed incomplete penetrance and highly variable age of disease onset (*4*). This variant, combined with the strong negative selection against both missense and protein-null variants in human population sequencing data, justified an extensive assessment of whether simple protein-null haploinsufficiency can cause neuropathy. We demonstrated that heterozygous knockout mice (with half of α1 subunit in their nerves, Figure 1), lack neuromotor deficiencies up to 18 months of age (equivalent to >56 years old in humans, www.jax.org) and are indistinguishable from wildtype mice. We compared ∼30 mice for each genotype at age 6 months (including exercise group) and ∼15 at 18 months and our data indicates that if lack of an α1 allele causes neuropathy, the disease penetrance would be very low and its age of onset would be very high. Furthermore, the comparable compound muscular action potential amplitude (Figure 5) suggests that these mice lack a subclinical disease. Thus, we conclude that the heterozygous *Atp1a1*^*+/-*^ knockout is <u>not</u> a model for *ATP1A1*-related CMT disease.

### Lack of effect of exercise on disease penetrance

Lifestyle could be a factor increasing the likelihood of CMT appearance in humans with *ATP1A1* mutations. A more active lifestyle with continuous exercise would normally be considered beneficial, but if the Na^+^/K^+^ pumping capacity is drastically altered, one could predict that increasing the number of action potentials propagating through the nerves would make the reduced pumping capacity in the axons more deleterious. In such a case, the Na^+^ overload could cause a rise in intracellular Ca^2+^ (due to the reduced driving force for the Na^+^/Ca^2+^ exchanger) and increased axonal death. Whether lifestyle differences may help explain the differences in family members with or without CMT remains unclear, but our results show that long term exercise did not cause CMT appearance by 6 months old in *Atp1a1*^*+/-*^ mice.

While not suggestive of a neuropathic disease state, it is interesting to note that the exercised wildtype cohort improved in its motor coordination (as evidenced by its increased latency to fall off of the rotarod compared to their non-exercised counterparts, Figure 5C). Here, we observed a median shift in latency to fall, to the maximum possible value, in exercised wildtype mice but not in *Atp1a1*^*+/-*^ mice. This indicates that exercise may result in contrasting differences between both genotypes and further study with greater numbers of mice and/or a more intensive accelerating rotarod protocol may be required for statistically significant differences to be observed. These possibilities will be evaluated in the future, with a modification of the standard rotarod protocol to discern beyond 180 s and/or with accelerations to faster speeds.

### A human research participant lacking one allele’s protein product is phenotypically normal

Consistent with the results in mice, we provide the first direct evidence for non-penetrance of heterozygous protein-null *ATP1A1* variants in humans. Deep phenotyping of our study participant with the p.(Y148*) variant confirmed the absence of clinical features of any of the known *ATP1A1*-related disorders into adulthood. While we did not directly measure transcript abundance *in vivo*, this variant is well before the last exon and predicted to cause nonsense-mediated decay, and could not be detected in membrane preparations of cells expressing a fluorescently tagged version of the early-termination mutant (Figure 6). We also cannot rule out the possibility that she could develop neuropathy in old age, or that the Raynaud phenomenon actually represents a small-fiber neuropathy which would be distinct from CMT and not detectable by NCS/EMG. Nevertheless, these results indicate that the spectrum of *ATP1A1* loss-of-function related phenotypes should be widened to include “healthy” individuals.

### Implications of incomplete penetrance of ATP1A1 protein-null variants

Taken together, our results in mice and a human raise the issue of whether missense variants might have different phenotypic severity from protein-null variants due to dominant-negative and/or compensatory mechanisms. Contrary to the expectation that haploinsufficiency would have similar or more deleterious effects than CMT2 missense alleles, which reduce Na^+^/K^+^ transport, we observed non-penetrance of protein-null alleles in both species.

In the first scenario, the presence of variant protein itself would be deleterious. Such dominant-negativity has been proposed for the paralogous *ATP1A3* gene, where pathogenic missense variants interfere with wildtype activity in the oocyte expression system (34). Furthermore, a human disease variant allele introduced into the nematode *eat-6* ortholog has a more severe phenotype than a whole-gene deletion allele in heterozygotes (*35*). Potential mechanisms of dominant-negativity of a nonfunctional alpha subunit include toxicity due to activation of the unfolded protein response (*36*) or competition for assembly with the beta subunit and trafficking to the cell membrane.

A second possibility is that activation of the genetic compensation response might decrease the severity of premature truncating variants compared to missense, as nonsense-mediated decay would trigger upregulation of transcription from both alleles, restoring wildtype mRNA expression (*37*). However, this cannot explain the lack of phenotype in the mice, which carried a premature truncating allele containing a frameshift deletion of exons 15-18 (corresponding to exons 17-20, c.2587-3142, of the NM_144900.2 reference transcript). Despite this, Atp1a1 protein levels remained at ∼50% with no obvious compensatory expression of α2/α3 paralogs.

One important limitation is that due to the extreme rarity of *ATP1A1* null variants, we cannot exclude the possibility of a low-penetrance phenotype. Given the strong selection against *ATP1A1* null variants in human sequencing datasets, it is still possible that some individuals with null alleles have severe or lethal disease that removes them from the pool of study participants. It is also possible that the higher attrition rate in *Atp1a1*^*+/-*^ relates to other pathological characteristics, yet to be identified. Phenotyping of additional patients and apparently “healthy” individuals is needed to establish the true spectrum of *ATP1A1-*related disease.

## Methods

### Animals

Heterozygous *Atp1a1* knockout (*Atp1a1*^*+/-*^) mice were obtained via a Material Transfer Agreement from the lab of Jerry Lingrel at the University of Cincinnati and delivered to the Lab Animal Resources Center (LARC) at Texas Tech University Health Sciences Center (TTUHSC). Animals used in the experimental procedures were derived using brother/sister mating from these progenitors. All mice were provided *ad libitum* access to food (LabDiet® 5R53) and filtered tap water. Animals were housed in Allentown IVC cages (NexGen™ Mouse 500) with 55 air changes per hour. Within these cages, animals were housed in groups of 2 to 5 with the exception that 5 cages of male mice required separation later in adulthood due to escalating aggression. Animals were maintained under a 12-hour light/dark cycle with humidity kept at 70%. All experimental procedures were approved by the Institutional Animal Care and Use Committee at TTUHSC. We studied both male and female mice and at least one exemplar per genotype per litter were studied to minimize confounding litter effects. Unexpected attrition due to spontaneous death was observed in five *Atp1a1*^*+/-*^ mice, in our experimental protocol (at 1, 5 (2 mice), 16 and 19 months of age). In contrast, no attrition was observed in wildtype mice, who always died by euthanasia.

### COVID-19 Statement

Establishment and maintenance of our breeding and experimental colony was significantly impacted by the COVID-19 pandemic. Funding for this mouse study begun in February 2020 and closure of the laboratory facilities to only essential staff early on in the pandemic precluded assessment of some cages of experimental animals at different ages. As a consequence, our planned longitudinal assessment of *Atp1a1*^*+/-*^ mice was stymied and we instead performed a cross-sectional evaluation of age-related deficits in *Atp1a1*^*+/*^ mice, while still maintaining strict comparison of littermates.

### Genotyping

Genotyping was performed at weaning from DNA isolated by either tail snip or ear punch. Platinum Direct PCR Universal Master Mix (ThermoFisher, A44647500) was used for DNA isolation and PCR. The recommended thermocycler protocol included with the kit was used. The following primers were ordered from IDT; wildtype allele forward and reverse primers: 5′-ATCAGAGCCAACAATCCC-3′ and 5′-GGTTAAATGGGAGGAGAATATG-3′ at a final concentration of 0.7 µM; knockout allele forward and reverse primers: 5′-TGGATCATACCTAAGTTGG-3′ and 5′-TACTCCCATTTGTCACGTC-3′ at a final concentration of 5 μM. Amplification of knockout and/or wildtype bands was identified via gel electrophoresis.

### Membrane Preparation and Western Blotting

Both sciatic nerves were dissected from five WT and six *Atp1a1*^*+/-*^ mice, pooled, and homogenized manually using a microcentrifuge pestle and then briefly sonicated in an NP-40 lysis buffer (150 mM NaCl, 1% NP-40 v/v, 50 mM Tris-Cl pH 8.0 with protease inhibitor cocktail (Sigma-Aldrich) and 1 mM PMSF. The homogenate was agitated for 2 hours at 4 °C, spun down at 12,000 rpm for 20 minutes and the supernatant collected for Western blotting. Protein concentration was determined by BCA assay (Thermo Fisher). Samples were incubated at 37 °C for 30 minutes in Laemmli buffer (to enhance denaturation and prevent aggregation) and loaded onto 7.5% mini-Protean TGX pre-cast gels (BioRad). After 45 min of electrophoresis (∼20 min at 90 V and ∼25 at 150 V) the protein was transferred to a polyvinylidene difluoride (PVDF) membrane by electroblot (30 mA for 3 hours). The PVDF membrane was blocked in PBS containing 0.1% v/v Tween-20 and 5% w/v milk for one hour. The membrane was incubated overnight at 4 °C with rabbit and mouse primary antibodies at 1:1000 dilutions; the polyclonal anti-α1 NASE or anti-α2 HERED (*32, a gift form Sandrine Pierre, Marshal Universtiy*) and the monoclonal anti-α3 XVIF9-G10 (Santacruz Biotech). The membrane was washed three times for 5-15 minutes in PBS with 0.1% v/v Tween-20 and incubated for one hour at room temperature in goat raised anti-rabbit (IRDye 680RD) and anti-mouse (IRDye 800CW) secondaries (LI-COR, 1:10,000). The membrane was washed three times in PBS with 0.1% Tween-20 and imaged using a ChemiDoc Imaging System (Bio-Rad). Plasma membrane enriched oocyte membrane protein was prepared as described (*38*).

### Mouse Behavioral Testing

Animals were tested on a series of validated behavioral paradigms associated with balance, strength, dexterity, motor coordination. Before testing, animals were moved to the LARC behavioral testing room, singly housed with food and water, and habituated to the test room conditions for at least 20 minutes. For the balance beam, pole test and accelerating rotarod animals were first trained over two days, three trials per day with an inter-trial interval of 1 minute, at one month of age. For testing in subsequent months, refamiliarization was performed only on a single day, with three trials and an inter-trial interval of 1 minute. The test session occurred on the day subsequent to either the last training or refamiliarization session with three trials and an inter-trial interval of 1 minute.

In the balance beam test, each animal walked across a white acrylic bar, 100 × 0.3 × 2.5 cm; L x W x D, with a start and end zone of 10 cm, respectively; each animal traversed a distance of 80 cm per trial. Our pole test utilized a PVC pole, 9 mm in diameter and 60 cm high with a start zone of 5 cm, and a test zone, covered with 250 grit sandpaper to a height of 55 cm. Animals were placed with the snout facing downwards and then trained to descend the pole. The balance beam and pole test test-sessions were video recorded using high-resolution cameras. In the accelerating rotarod, animals walked/ran on an accelerating motor driven drum with an initial speed of 4 revolutions per minute (RPM) and a final speed of 40 RPM. Each trial took a maximum of 3 minutes. All data included here were obtained with a six lane rotarod from Maze Engineers with are 3 cm diameter rod. The primary outcome variable was the latency to fall off of the rod. As previously described (*39-41*), the behavior where a mouse simply grasped the groves in the bar and rotated around the bar, rather than posturally adjusting to walk/run on the bar, was equivalent to falling. Initially, we performed experiments using a Rotamex 4/8 apparatus with a 3.875 cm drum where we tested four animals, two of each genotype, simultaneously. After several tests, a mechanical error developed in the apparatus, where we noticed that animals were falling more quickly in the first and last lane than in the two middle lanes. The difference in drum size of the two apparatus makes it difficult to compare, and only the data from the Maze Engineers rotarod were included.

In the hindlimb clasping test, animals were suspended by the tail and video recorded for a minimum of twelve seconds. Consistent with the original protocol only the first twelve seconds of video were scored (*21-23*). In our administration of this test, we discovered that small methodological differences, not reported previously, significantly impacted the degree of splaying observed at the individual mouse level. If the mouse was grasped by the base of the tail (close to the anus), no or little splaying of the hindlimbs was observed. Moving from the base of the tail to the tip of the tail induced more splaying, while grasping the animal in the middle of the tail induced some splaying. Moreover, the proximity of the animal’s forelimbs to a solid substrate was sufficient to affect splaying, with greater splaying being observed when the animal was further from the substrate (such as a cage lid) and less splaying when the animal was closer to or grasping the substrate. Because of these previously unreported factors that affected the outcome being measured, we formalized a final protocol where the mice were hung by the tip of the tail, suspended over an empty cage with a grid lid, with the forelimbs approximately 3 cm from the lid. Only these data are described here.

### Electromyography

CMAP was measured in isoflurane anesthetized mice (4% for induction and 2% for maintenance using a Sierra Summit EMG (Cadwell) at 18 months old. The mouse temperature was held constant at 37 °C by a heating pad and an anal temperature probe (Kent Scientific). The hindlimbs of the mice were shaved and defolliculated to insert disposable sterile subdermal 13 mm needle electrodes (Friendship Medical). The recording electrode was placed in the footpad, the ground in the intrinsic muscles of the back above the level of the sciatic nerve, the reference next to the Achilles tendon and the stimulating electrode in the sciatic notch. The stimulus was increased until the response amplitude reached a supramaximal stimulus. The CMAP amplitude was the difference between the maximum positive and negative peaks.

### Aldosterone Serum Analysis

The mice were anesthetized via isoflurane and the abdomen was opened to access the thoracic cavity and pericardial cavity. Cardiac puncture of the right ventricle with a 23-gauge needle was used to collect blood samples. The whole blood was allowed to clot and spun down to isolate serum. Serum aldosterone concentration was determined by ELISA following the manufacturer protocol (Aldosterone ELISA kit, Enzo Life Sciences).

### Statistical Analysis

Due to a high degree of skewness in some of our measures, as well as violation of the assumption of homogeneity of variance, our behavioral data were analyzed only using non-parametric techniques. Each month was analyzed separately and differences between the sexes and genotype were first evaluated via the Kurskall-Wallis H-test followed by Dunn’s corrected pair-wise comparisons. In this analysis, sex differences in general were observed but not with respect to an interaction with genotype. The observed sex differences likely relate to overall differences in body weight and agility that is typical of male and female mice (*18*). Consequently, we combined the data from both sexes and re-analyzed the data using the Mann Whitney U-Test.

### Clinical phenotyping

Human subjects provided written informed consent for study participation and publication under National Institutes of Health (NIH) Institutional Review Board protocol 76-HG-0238 (ClinicalTrials.gov NCT00369421). The p.(Y148*) variant was identified on research genome sequencing of the proband by the NIH Reverse Phenotyping Core (ClinicalTrials.gov NCT03632239), a population sequencing study in which participants could be re-contacted. Deep phenotyping was performed by board-certified neurologists and medical geneticists at the NIH Clinical Center, including neurological exam, blood and urine sampling, and nerve conduction study/electromyography (NCS/EMG).

### Molecular Biology and HEK Cellular Culture

The full-length wildtype cDNA for the YFP N-terminally conjugated α1 and the β1 subunits (a gift from Seth L. Robia at Loyola University Chicago) have been previously described (*42*). The Y148* point variant was introduced via site-directed mutagenesis PCR and verified by sequencing.

Human embryonic kidney cells (HEK 293) were grown in Dulbecco’s modified Eagle’s medium (Gibco) with 10% v/v fetal bovine serum (Gibco) and 2% v/v penicillin/streptomycin (Gibco) and incubated at 37°C and 5% CO_2_ in a humidified incubator (Thermo Scientific). At 60-80% confluency the cells were transiently transfected (Lipofectamine 3000 kit; Thermo Fisher) with α (either wildtype or Y148*) and β subunit cDNA in a 3:1 ratio. Forty-eight hours after transfection cells were scraped to make the membrane preparation (see next section). A few scraped cells were replated at a lower density on poly-D-lysine coated coverslips (Electron Microscopy Sciences) for imaging following fixation 24 hours after replating.

### HEK cell membrane preparation

After 48 hours the transfected cells were washed with 1XPBS and cellular homogenizing buffer (250mM sucrose, 10mM Tris, and 2 mM EDTA, 1mM PMSF, and 1x protease inhibitor cocktail (Sigma)). The cells were scraped on ice, collected, pelleted, and homogenized using a microcentrifuge dounce. The homogenate was centrifuged at 4,000g for 20 min and the resulting supernatant was collected. The pellet was resuspended and centrifuged again at 4,000g for 20 minutes and combined with the supernatant from the first centrifuge step. The combined supernatant was centrifuged at 55,000g for 30 min. The membrane fraction pellet was resuspended in homogenizing buffer. Protein quantification and western blotting was carried out as described above. After transfer and blocking, the PVDF membrane was probed with primary antibodies rabbit anti-NASE (1:1000) and mouse anti-YFP (1:1000; N86/8, DSHB).

### Confocal Imaging

Twenty-four hours after replating, the coverslips were washed in 1X PBS and fixed with 4% v/v paraformaldehyde (MP Biomedicals) for 10 minutes. Coverslips were mounted (ProLong Diamond Antifade Mountant, Thermo Fisher) and imaged using Nikon T1-E microscope with A1 confocal (TTUHSC Image Analysis Core Facility).

## Acknowledgements

We thank Sandrine Pierre and the late Jerry B. Lingrel for supplying the *Atp1a1*^*+/-*^ colony. This work was supported by grants from NINDS 1-R03 NS116433-01 to PA and JDB and supplement 3R03NS116433-02s1 to PA and AP, and by NHLBI 1R01HL158649-01A1 to Seth L. Robia and PA. MH, AK, FR, CT, CW, and SY were supported by the NIH Intramural Research Program. The members of the Reverse Phenotyping Core are as follows: Clesson Turner, Alexander E. Katz, Caralynn M. Wilczewski, Felicia Akinwande, Vera Zanker, Tyra G. Wolfsberg, Suiyuan Zhang, Sumeeta Singh, Justin E. Paschall, George L. Maxwell, Morgan Similuk, and Leslie G. Biesecker.

## Notes

### Competing Interest Statement

The authors have declared no competing interest.

## References

1. F. Beuschlein et al., Somatic mutations in ATP1A1 and ATP2B3 lead to aldosterone-producing adenomas and secondary hypertension. Nature genetics 45, 440–444, 444e441-442 (2013).

2. E. A. Azizan et al., Somatic mutations in ATP1A1 and CACNA1D underlie a common subtype of adrenal hypertension. Nature genetics 45, 1055-1060 (2013).

3. T. A. Williams et al., Somatic ATP1A1, ATP2B3, and KCNJ5 Mutations in Aldosterone-Producing Adenomas. Hypertension 63, 188–195 (2014).

4. P. Lassuthova et al., Mutations in ATP1A1 Cause Dominant Charcot-Marie-Tooth Type 2. Am J Hum Genet 102, 505–514 (2018).

5. J. He et al., ATP1A1 mutations cause intermediate Charcot-Marie-Tooth disease. Hum Mutat 40, 2334–2343 (2019).

6. F. Stregapede et al., Hereditary spastic paraplegia is a novel phenotype for germline de novo ATP1A1 mutation. Clin Genet 97, 521–526 (2020).

7. K. P. Schlingmann et al., Germline De Novo Mutations in ATP1A1 Cause Renal Hypomagnesemia, Refractory Seizures, and Intellectual Disability. Am J Hum Genet 103, 808–816 (2018).

8. E. D. Biondo, K. Spontarelli, G. Ababioh, L. Mendez, P. Artigas, Diseases caused by mutations in the Na(+)/K(+) pump alpha1 gene ATP1A1. Am J Physiol Cell Physiol 321, C394–C408 (2021).

9. F. Cinarli Yuksel et al., The phenotypic spectrum of pathogenic ATP1A1 variants expands: the novel p.P600R substitution causes demyelinating Charcot–Marie–Tooth disease. Journal of Neurology, (2023).

10. D. J. Meyer, C. Gatto, P. Artigas, Na/K Pump Mutations Associated with Primary Hyperaldosteronism Cause Loss of Function. Biochemistry 58, 1774–1785 (2019).

11. D. J. Meyer, C. Gatto, P. Artigas, On the effect of hyperaldosteronism-inducing mutations in Na/K pumps. J Gen Physiol 149, 1009-1028 (2017).

12. S. Ygberg et al., A missense mutation converts the Na(+),K(+)-ATPase into an ion channel and causes therapy-resistant epilepsy. J Biol Chem 297, 101355 (2021).

13. K. J. Karczewski et al., The mutational constraint spectrum quantified from variation in 141,456 humans. Nature 581, 434–443 (2020).

14. L. C. Kutz et al., Isoform-specific role of Na/K-ATPase alpha1 in skeletal muscle. Am J Physiol Endocrinol Metab 314, E620–E629 (2018).

15. L. C. Barcroft, A. E. Moseley, J. B. Lingrel, A. J. Watson, Deletion of the Na/K-ATPase alpha1- subunit gene (Atp1a1) does not prevent cavitation of the preimplantation mouse embryo. Mech Dev 121, 417–426 (2004).

16. A. E. Moseley et al., Genetic profiling reveals global changes in multiple biological pathways in the hearts of Na, K-ATPase alpha 1 isoform haploinsufficient mice. Cell Physiol Biochem 15, 145–158 (2005).

17. R. J. Carter, J. Morton, S. B. Dunnett, Motor coordination and balance in rodents. Curr Protoc Neurosci Chapter 8, Unit 8 12 (2001).

18. D. Wahlsten, Mouse Behavioral Testing: How to Use Mice in Behavioral Neuroscience. (Elsevier Science, 2010).

19. Y. Ohno, Ishida, K., Ieda, K., Ishibashi, T., Okada, K., Nakamura, M., Evaluation of bradykinesia induction by SM-9018, a novel 5-HT2 and D2 receptor antagonist, using the mouse pole test. Pharmacology, Biochemistry, & Behavior 49, 19–23 (1994).

20. N. Ogawa, Y. Hirose, S. Ohara, T. Ono, Y. Watanabe, A simple quantitative bradykinesia test in MPTP-treated mice. Res Commun Chem Pathol Pharmacol 50, 435–441 (1985).

21. J. Lee et al., Overexpression of mutant HSP27 causes axonal neuropathy in mice. J Biomed Sci 22, 43 (2015).

22. T. T. Sabblah et al., A novel mouse model carrying a human cytoplasmic dynein mutation shows motor behavior deficits consistent with Charcot-Marie-Tooth type 2O disease. Sci Rep 8, 1739 (2018).

23. R. de Haas, F. G. Russel, J. A. Smeitink, Gait analysis in a mouse model resembling Leigh disease. Behav Brain Res 296, 191–198 (2016).

24. L. L. Teunissen, Disease Course of Charcot-Marie-Tooth Disease Type 2. Archives of Neurology 60, 823–828 (2003).

25. M. Bohlen et al., Experimenter effects on behavioral test scores of eight inbred mouse strains under the influence of ethanol. Behav Brain Res 272, 46–54 (2014).

26. B. Martin, S. Ji, S. Maudsley, M. P. Mattson, “Control” laboratory rodents are metabolically morbid: why it matters. Proc Natl Acad Sci U S A 107, 6127-6133 (2010).

27. J. D. Bailoo et al., Effects of Cage Enrichment on Behavior, Welfare and Outcome Variability in Female Mice. Front Behav Neurosci 12, 232 (2018).

28. J. D. Bailoo et al., Evaluation of the effects of space allowance on measures of animal welfare in laboratory mice. Sci Rep 8, 713 (2018).

29. J. D. Bailoo et al., Effects of weaning age and housing conditions on phenotypic differences in mice. Sci Rep 10, 11684 (2020).

30. J. H. Kaplan, Biochemistry of Na,K-ATPase. Annu Rev Biochem 71, 511–535 (2002).

31. C. Gatto, S. M. McLoud, J. H. Kaplan, Heterologous expression of Na+-K+-ATPase in insect cells: intracellular distribution of pump subunits. American Journal of Physiology - Cell Physiology 281, 982–992 (2001).

32. T. A. Pressley, Phylogenetic conservation of isoform-specific regions within alpha-subunit of Na(+)-K(+)-ATPase. Am J Physiol 262, C743–751 (1992).

33. P. Rigoard et al., The Na, K-ATPase alpha3-isoform specifically localizes in the Schmidt-Lanterman incisures of human nerve. Cellular and Molecular Biology 53, 1003–1009 (2007).

34. M. Li et al., A functional correlate of severity in alternating hemiplegia of childhood. Neurobiol Dis 77, 88–93 (2015).

35. A. Sorkaç, I. C. Alcantara, A. C. Hart, In Vivo Modelling of ATP1A3 G316S-Induced Ataxia in C. elegans Using CRISPR/Cas9-Mediated Homologous Recombination Reveals Dominant Loss of Function Defects. PloS one 11, e0167963 (2016).

36. E. Arystarkhova, L. J. Ozelius, A. Brashear, K. J. Sweadner, Misfolding, altered membrane distributions, and the unfolded protein response contribute to pathogenicity differences in Na,K- ATPase ATP1A3 mutations. J Biol Chem 296, 100019 (2021).

37. Z. Ma et al., PTC-bearing mRNA elicits a genetic compensation response via Upf3a and COMPASS components. Nature 568, 259–263 (2019).

38. D. J. Meyer et al., FXYD protein isoforms differentially modulate human Na/K pump function. Journal of General Physiology 152, e202012660 (2020).

39. B. J. Jones, & Roberts, D. J., The quantitative measurement of motor inco-ordination in naive mice using an accelerating rotarod. Journal of Pharmacy and Pharmacology 4, 302–304 (1968).

40. N. W. Dunham, T. S. Miya, A note on a simple apparatus for detecting neurological deficit in rats and mice. J Am Pharm Assoc Am Pharm Assoc 46, 208–209 (1957).

41. M. O. Bohlen, J. D. Bailoo, R. L. Jordan, D. Wahlsten, Hippocampal commissure defects in crosses of four inbred mouse strains with absent corpus callosum. Genes Brain Behav 11, 757–766 (2012).

42. J. Seflova et al., Fluorescence lifetime imaging microscopy reveals sodium pump dimers in live cells. J Biol Chem 298, 101865 (2022).

